# *Verticillium dahliae* inoculation and *in vitro* propagation modify the xylem microbiome and disease reaction to Verticillium wilt in a wild olive genotype

**DOI:** 10.1101/2020.11.23.392977

**Authors:** Manuel Anguita-Maeso, José Luis Trapero, Concepción Olivares-García, David Ruano-Rosa, Elena Palomo-Ríos, Rafael M. Jiménez-Díaz, Juan A. Navas-Cortés, Blanca B. Landa

## Abstract

Host resistance is the most practical, long-term and economically efficient disease control measure for Verticillium wilt in olive caused by the xylem-invading fungus *Verticillium dahliae* (*Vd*), and it is at the core of the integrated disease management. Plant’s microbiome at the site of infection may have an influence on the host reaction to pathogens; however, the role of xylem microbial communities in the olive resistance to *Vd* has been overlooked and remain unexplored to date. This research was focused on elucidating whether *in vitro* olive propagation may alter the diversity and composition of the xylem-inhabiting microbiome and if those changes may modify the resistance response that a wild olive clone shows to the highly virulent defoliating (D) pathotype of *Vd*. Results indicated that although there were differences in microbial communities among the different propagation methodologies, most substantial changes occurred when plants were inoculated with *Vd*, regardless whether the infection process took place, with a significant increase in the diversity of bacterial communities when the pathogen was present in the soil. Furthermore, it was noticeable that olive plants multiplied under *in vitro* conditions developed a susceptible reaction to D *Vd*, characterized by severe wilting symptoms and 100% vascular infection. Moreover, those *in vitro* propagated plants showed an altered xylem microbiome with a decrease in total OTU numbers as compared to that of plants multiplied under non-aseptic conditions. Overall, 10 keystone bacterial genera were detected in olive xylem regardless infection by *Vd* and the propagation procedure of plants (*in vitro* vs nursery), with *Cutibacterium* (36.85%), *Pseudomonas* (20.93%), *Anoxybacillus* (6.28%), *Staphylococcus* (4.95%), *Methylobacterium-Methylorubrum* (3.91%), and *Bradyrhizobium* (3.54%) being the most abundant. *Pseudomonas* spp. appeared as the most predominant bacterial group in micropropagated plants and *Anoxybacillus* appeared as a keystone bacterium in *Vd*-inoculated plants irrespective of their propagation process. Our results are first to show a breakdown of resistance to *Vd* in a wild olive that potentially maybe related to a modification of its xylem microbiome, and will help to expand our knowledge of the role of indigenous xylem microbiome on host resistance which can be of use to fight against main vascular diseases of olive.

## Introduction

Verticillium wilt, caused by the vascular-colonizing, soil-borne fungus, *Verticillium dahliae* (*Vd*), is one of the main diseases threatening the health and growth of olive (*Olea europaea* subsp. *europaea* var. *europaea*) production worldwide. This disease, first reported in Italy in 1946, has steadily increase in prevalence and incidence to become an actual threat to olive cultivation in the Mediterranean Basin due to the high rates of tree mortality and important reductions in yields (Jiménez-Díaz et al., 2011; Landa et al., 2019). Infection of olive plants by *Vd* takes place through the root system; then the pathogen colonizes the xylem vessels and impairs the sap flow by means of mycelial proliferation, the formation of occlusions and tyloses that ultimately cause the tree death (Báidez et al., 2007; Jiménez-Díaz et al., 2011). Two pathotypes have been identified among *Vd* isolates in olive, namely defoliating (D) and non-defoliating (ND), which differ much in virulence and determine severity of Verticillium wilt. The ND pathotype induces moderately severe branch die back and leaf necrosis, whereas the highly virulent D pathotype induces severe falling of green leaves, necrosis of entire plant canopy sectors and eventually tree death (Navas-Cortés et al., 2008; Jiménez-Díaz et al., 2011). In Andalusia, the world’s largest producer of olive oil, circa 39% of old and newly planted olive orchards were reported affected by Verticillium wilt, but inspections of 90 arbitrarily chosen orchards in the three main olive-growing provinces of the region indicated 71% disease prevalence with a mean incidence of 20% in the affected orchards (Jiménez-Díaz et al., 2011).

The most practical and economically efficient method for the management of Verticillium wilt is the use of resistant cultivars. However, most of olive cultivars widely grown in Spain, are moderately to highly susceptible to D *Vd* (López-Escudero et al., 2010; Jiménez-Díaz et al., 2011; Ostos et al., 2020). Recently, a few wild olive genotypes were identified as highly resistant to D *Vd* that may have a valuable potential as rootstocks for the management of Verticillium wilt (Jiménez-Fernández et al., 2016). Nevertheless, use of a single control measure may not be fully effective for the management of Verticillium wilt in olives, as shown for other wilt diseases. Thus, an integrated management strategy it is advisable, combining the use of resistant olive cultivars, or of tolerant ones grafted onto resistant rootstocks, with adequate irrigation management and agricultural practices that prevent the spread of inoculum of the pathogen (Jiménez-Díaz et al., 2011; Jiménez-Fernández et al., 2016).

Although plants have evolved their own adaptations to alleviate most biotic and abiotic stresses in nature, they also rely on their microbial partners to survive and defend themselves against microbial invaders and pathogens (Hassani et al., 2018). In nature, healthy plants live in association and interact with a myriad of microorganisms, collectively called the plant microbiome, which is now known to bear important roles in plant health. Endophytes are bacteria and fungi that live within plants where they establish nonpathogenic relationships with their hosts (Azevedo et al., 2000). Endophytic microbial communities are dynamic and are influenced by several factors. These communities promote plant growth directly by phytostimulation and biofertilization, and/or indirectly by inducing stress tolerance and disease suppression (Compant et al., 2016; Hassani et al., 2018). Therefore, a thorough knowledge of the microbial communities residing within the xylem vessels of olive trees may be crucial for understanding their potential influence on the healthy growth of this plant as well as on the resistance shown by specific olive genotypes against D *Vd* or other vascular plant pathogens (Hong and Park, 2016). Thus, certain members of the microbial community may play a crucial role in the expression of disease resistance in plants to plant pathogens by setting on diverse mechanisms that mainly include a via for nutrient mobilization, direct antagonism, niche exclusion and induction of systemic and localized resistance (Müller et al., 2015; Hassani et al., 2018; Wille et al., 2019). However, although it can be considered that most of the olive xylem microbiota should be assembled by microorganisms with neutral or positive effects, their mechanistic role in defense against vascular pathogens has not yet been addressed.

Knowledge of the olive tree xylem microbiome is limited and very fragmentary, despite progress made on understanding the structure and functions of the plant microbiome in recent years. Possibly, technical difficulties for isolation of xylem microbiome have difficulted its study, and only a few have focused on microorganisms inhabiting the olive xylem. However, different methodological approaches, including culture-dependent procedures complemented with next-generation sequencing (NGS) technologies, have recently made possible to characterize microbial communities associated with olive xylem tissue (Hardoim et al., 2015; Fausto et al., 2018; Anguita-Maeso et al., 2020; Zicca et al., 2020).

The modification or attenuation of the diversity and composition of xylem microbial communities might result in different responses from the plant host to cope with vascular pathogens. The transmission of microbiota to the progeny in plants vegetatively propagated represents a way to ensure the presence of beneficial symbionts within the plant (Vannier et al., 2018; Liu et al., 2019). However, it is unknown how xylem endophytic microbiota in olive may be transmitted from shoot tips to explants, as well as to mature plants, and how stable would that be during this process. Micropropagation has become an important tool to reproduce selected olive genotypes and guarantees true to type and pathogen free plants (Fabbri et al., 2009). Olive micropropagation, through tissue culture, which is conducted *in vitro* under aseptic conditions, for at least a certain period of time, can potentially induce changes an alter the composition of the xylem microbiome. Thus, by producing olive plants by tissue culture, some beneficial and non-pathogenic endophytes may be excluded, since tissue cultures are initiated from shoots after extensive surface sterilization and then plants are maintained under aseptic conditions. Consequently, this propagation procedure represents an ideal experimental approach to test whether the xylem microbiome has a role on the resistance shown by specific olive genotypes to vascular pathogens.

This present research was conceived to elucidate how plants protect themselves by shaping their xylem microbiome in a resistant wild olive genotype as a first step to assess the complex plant–microbe interactions in the xylem that can contribute to maintain olive health and its resistance response against vascular pathogens. The specific objectives of this work were to determine whether: i) *in vitro* propagation methodology can modify the diversity and composition of the xylem microbiome in olive; ii) those changes may alter host resistance response to the vascular pathogen *Vd;* and iii) inoculation and vascular infection of olive by *Vd* may induce changes in the olive xylem microbiome. Determining those effects may be essential to the identify microbial communities that are triggered after pathogen infection and could act as antagonists against *Vd*. Furthermore, understanding the tight relationships between xylem-inhabiting microorganisms and vascular pathogens will help to reveal determinant microbial players that may contribute to produce olive plants more resilient to infection by vascular pathogens.

## Material and methods

### Olive Plant material

A wild olive (*Olea europaea* var. *sylvestris*) clone ‘Ac-18’ highly resistant to the D pathotype of *Vd* (Jiménez-Fernández et al., 2016) was used in this study. The highly resistance of ‘Ac-18’ plants to D *Vd* is characterized by symptomless infection, together with plugging of xylem vessels, no re-isolation of the fungus from stem vascular tissue and the plant’s ability to quantitatively reduce the extent of stem colonization by the pathogen (Jiménez-Fernández et al., 2016). Also, olive cv. ‘Picual’ highly susceptible to D and susceptible to ND *Vd* pathotypes (Calderon et al., 2014), and a wild olive clone ‘Ac-15’ highly susceptible to D *Vd* (Narváez et al., 2019) were used in the pathogenicity experiments as controls of disease reaction.

Three types of ‘Ac-18’ plant material were used in the study, which were derived from shoots of a same mother adult plant, named: i) *in vitro*-standard plants: plants micropropagated using the standard olive methodology of axillary shoot elongation; ii) *in vitro*-adapted plants: plants micropropagated and subsequently adapted to greenhouse conditions; and iii) nursery propagated: plants propagated following standard semi-woody stacking procedure at a commercial olive nursery.

For *in vitro*-standard propagation of ‘Ac-18’ and ‘Ac-15’ plants, 1.5 to 2.0 cm long shoots bearing two nodes were multiplied as in Narváez et al. (2020) using RP medium [DKW macro- and micronutrients and vitamins as modified by Roussos and Pontikis (2002)] supplemented with 2 mg/L zeatin riboside (Vidoy-Mercado et al., 2012). For rooting, 2-cm long shoots were cultured for 3 days in basal RP liquid medium supplemented with 10 mg/L IBA and subsequently transferred to basal solid RP medium supplemented with 1 g/L activated charcoal. Plants were incubated under a 16 h photoperiod at 40 μmol m^-2^ s^-1^ at 25±2°C for 8 weeks until ensuring at least three to four roots. Adaptation to *ex vitro* conditions was carried out initially using 35×24×18 cm germination boxes under high levels of sterility. Propagated ‘Ac-18’ rootlets were transplanted to 300 ml pots filled with a sterilized (121°C for 20 min) perlite and vermiculite (1:1) mixture. Plants were sprayed with sterile water, and the cover was sealed with transparent film, and incubated at 25±2°C in darkness in a growth chamber. After 2 days, a 2-h cycle of indirect fluorescent light of 360 μmol m^-2^ s^-1^ was provided, which duration was increased daily until reaching a 12-h light cycle within 1 week. The starting high relative humidity provided by the closed environment made unnecessary watering the plants for 3 weeks. Afterwards, plants were watered (3 ml per pot) weekly with sterile water using a 10-ml syringe and fertilized once per month with Hoagland’s nutrient solution (Hoagland and Arnon, 1950). After 8 weeks, plants were transplanted to 500 ml pots filled with a sterile peat:perlite (3:1) mixture. Plants grew for additional 6 months in a growth chamber adjusted to 22±2°C, 60–80% relative humidity and a 14 h photoperiod of fluorescent light of 360 μmol m^-2^ s^-1^. Plants were watered as needed and fertilized weekly with 100 mL Hoagland’s nutrient solution.

*In vitro*-adapted ‘Ac-18’ plants were produced using the same methodology than for *in vitro*-standard plants, with the exception that plants were grown for additional 12 months in a glass greenhouse at a fluctuating minimum/maximum temperature of 15±5 and 25±5°C across the entire growing period and daylight conditions. Plants were watered using tap water as needed and fertilized weekly as described before.

Finally, nursery propagated ‘Ac-18’ and ‘Picual’ plants followed standard semi-woody stacking procedure at a commercial olive nursery (Plantas Continental S.A, Córdoba, Spain). Briefly, semi-hard stem cuttings with two active leaves on the top were dipped in an indole butyric acid solution to stimulate rooting and planted on peat:coconut fiber (1:5) pellets under mist conditions in plastic tunnels (Caballero and Del Río, 2010). Once callus was formed and roots appeared (three to four roots) plants were transplanted to 500 ml pots containing a perlite:coconut fiber:peat (1.5:5:3.5) mixture amended with 4 g l^-1^ of slow release fertilizer (Osmocote^®^ Exact standard 15-9-12+2MgO; ICL Specialty Fertilizers, The Netherlands). Plants were incubated under natural environmental conditions for 6 months in a plastic greenhouse. During this time, plants received water as needed but no additional fertilizers.

At the time of inoculation with the pathogen *in vitro*-standard, nursery propagated and *in vitro-adapted* ‘Ac-18’ plants were 10-, 10- and 18-months old, respectively. Apparently, plants from all type of propagation procedures had a similar degree of bark lignification of the main stem and root development (**Supplementary Fig. 1**).

### Pathogenicity experiment

A monoconidial *Vd* isolate (V138I) from defoliated ‘Coker 310’ cotton plants at Córdoba (Spain), and representative of the D pathotype was used in the experiment. This isolate proved highly virulent on olive in previous work (Jiménez-Fernández et al., 2016). Inoculum consisted of an infested cornmeal-sand mixture (CMS; sand:cornmeal:deionized water, 9:1:2, w/w) produced as described by Jiménez-Fernández et al. (2016). The infested CMS was homogenized, allowed to desiccate in an incubator adjusted to 33°C for 3 days, and thoroughly mixed with a pasteurized soil mixture (clay loam:peat, 2:1, v/v) at a rate of approximately 1:20 (w/w) to reach an inoculum density of 5 × 10^7^ CFU g^-1^ soil of *Vd* as determined by dilution-plating on chlortetracycline-amended water agar (CWA; 1 L distilled water, 20 g agar, 30 mg chlortetracycline) (Jiménez-Fernández et al., 2016).

Plants of ‘Ac-18’ clone, representative of each propagation procedure, and susceptible ‘Picual’ and ‘Ac-15’ plants, were then transplanted to 1500 ml pots filled with the D *Vd*-infested soil mixture. Before transplanting, plants were uprooted from the potting substrate, gently shaken to retain only the rhizosphere soil and placed in pots filled with the infested soil mixture. Non-inoculated plants were treated similarly and transplanted to the pasteurized soil mixture mixed with non-infested CMS at the same rate as infested CMS. Inoculated and control plants were incubated in a growth chamber adjusted to 22±2°C, 60–80% relative humidity and a 14-h photoperiod of fluorescent light of 360 μmol m^-2^ s^-1^ for 3 months. During this time, plants were watered as needed with tap water and fertilized weekly as previously described. There were six replicated pots (one plant per pot) for inoculated and non-inoculated plants of each plant genotype, respectively, in a completely randomized design.

Disease reaction was assessed by the incidence (percentage of plants showing disease symptoms) and severity of foliar symptoms. Symptoms were assessed on individual plants on a 0 to 4 rating scale according to the percentage of affected leaves and twigs at 2- to 3-day intervals throughout the duration of the experiment (Jiménez-Fernández et al., 2016). Upon termination of the experiment, the extent of colonization by *Vd* was determined by isolations of the fungus in CWA (Jiménez-Fernández et al., 2016) from 6-cm-long stem pieces sampled from the main stem at the same time than similar samples were processed for extraction of xylem microbiome (see below). Data of pathogen isolation from the stem were used to calculate the intensity of stem vascular colonization for each individual plant, according to a stem colonization index (SCI) as described before (Jiménez-Fernandez et al., 2016).

Additionally, the amount of *Vd* present in ‘Ac-18’ stem samples was determined by using the TaqMan qPCR assay developed by Bilodeau et al. (2012) as described in Gramaje et al. (2013). The same DNA samples used for the xylem microbiome characterization were used for pathogen quantification, with each sample being analyzed in duplicate. All qPCR assays were performed in a LightCycler480 (Roche Diagnostics) apparatus. The cycle threshold (Ct) values for each qPCR reaction was calculated using the default estimation criteria in the manufacturer’s software. The quantification limit of the TaqMan qPCR assay was fixed at a Ct of 36 (0.1 pg of *Vd* DNA μl^-1^) (Gramaje et al., 2013).

### DNA xylem microbiome extraction and sequencing

The xylem microbiome was extracted following the procedure described by Anguita-Maeso et al. (2020). Briefly, a number of three 6-cm-long pieces from the bottom, middle and upper stem of each ‘Ac-18’ plant were debarked and xylem chips were obtained by scraping the most external layer of the debarked woody pieces with a sterile scalpel. Xylem chips from a ‘Ac-18’ plant were mixed together, and a 0.5-g sample was placed in a Bioreba bag containing 5 ml of sterile phosphate-buffered saline (PBS), the bags were closed with a thermal sealer and the content was macerated with a hand homogenizer (BIOREBA, Reinach, Switzerland). Extracts were stored at −80°C until DNA extraction. All the processes described above took place under sterile conditions within a flow hood chamber (Anguita-Maeso et al., 2020). Aliquots (0.5 ml) of macerated xylem chips in PowerBead tubes (DNeasy PowerSoil Kit, QIAGEN) were homogenized 7 min at 50 pulses s^-1^ with the Tissuelyser LT (QIAGEN) prior to incubation in lysis buffer for 1 h at 60°C for increasing cell lysis, and then processed following the DNeasy PowerSoil Kit manufacturer’s protocol.

Extracted DNA was used directly to amplify the V5-V6 rRNA region with the primers 799F (5’-AACMGGATTAGATACCCKG-3’) and 1115R (5’-AGGGTTGCGCTCGTTG-3’). PCR products were purified using Agencourt AMPure XP (Beckman Coulter) following manufacture instructions. Purified PCR products were quantified using Quant-iT™ PicoGreen™ dsDNA Assay Kit (Thermo Fisher Scientific) and a Tecan Safire microplate reader (Tecan Group, Männedorf, Switzerland). Equimolecular amounts from each individual sample in 10 mM of Tris were combined and the pooled library was sequenced by the Genomics Unit at ‘Fundación Parque Científico de Madrid’, Madrid, Spain using the Illumina MiSeq platform (Nano-V2; PE 2x 250 bp). The ZymoBIOMICS microbial standard (Zymo Research Corp., Irvine, CA, USA) and water (no template DNA) were used as internal positive and negative controls, respectively, for library construction and sequencing. Raw sequence data have been deposited in the Sequence Read Archive (SRA) database at the NCBI under BioProject accession number PRJNA679263.

### Statistical and bioinformatics analyses

Quality control and adapter trimming of demultiplexed raw fastq 16S rRNA sequences obtained from MiSeq output was performed with FastQC and TrimGalore tools. Truncation length in 225 bp of the forward and reverse reads was needed to increase the Phred quality (Q>30) score visualized in MultiQC tool. No trimming length was needed. Quality reads were then analyzed using the Quantitative Insights into Microbial Ecology bioinformatics pipeline, QIIME2 (version 2020.2; https://view.qiime2.org/) (Caporaso et al., 2010; Bolyen et al., 2019) with default parameters unless otherwise noted. DADA2 pipeline were used for denoising fastq paired-end sequences along with filtering chimeras. Operational taxonomic units (OTUs) were obtained at 99% of similarity and were taxonomically classified using VSEARCH consensus taxonomy classifier (Rognes et al., 2016) against Silva SSU v.138 database. Singletons were discarded for taxonomy and statistical analysis.

Differences in bacterial communities were calculated in QIIME2 using rarefaction curves of alpha-diversity indexes (Richness, Shannon, and Simpson) at OTU level. Alpha and beta diversity as well as alpha rarefaction curves were conducted rarefying all samples to the minimum number of reads found. The nonparametric Scheirer–Ray–Hare test (*P* <0.05) was used to assess the effects of the inoculation treatment, propagation method and their interaction in alpha diversity indexes, using the rcompanion package (Mangiafico, 2020) in R. Dunn’s Kruskal–Wallis multiple comparisons were performed for post hoc analysis. The *P*-value was adjusted with the Benjamini–Hochberg method (Benjamini and Hochberg, 1995). Venn diagrams were generated using Venn package (Dusa, 2018) in R were used to identify shared (core microbiome) or unique taxa according to the inoculation treatment and propagation methods. Linear discriminant analysis effect size (LEfSe) method to identify differentially abundant bacterial taxa associated to inoculation treatments and propagation methods (Segata et al., 2011).

Additionally, weighted UniFrac distances were estimated at OTU level taking into account the phylogenetic distance among bacterial communities (Lozupone and Knight, 2005). Principal coordinates analysis (PCoA) of weighted UniFrac distance matrix was used to evaluate similarities among the bacterial communities according to the inoculation treatment or propagation procedure. Additionally, the *adonis* function within the vegan package in R (999 permutations) was performed to test the effects (*P*< 0.05) of the inoculation treatment, the plant propagation method and their interaction.

## Results

### Pathogenicity experiment

Typical Verticillium wilt symptoms consisting on early dropping of green leaves, and necrosis and death of some branches, characteristics of infection by D *Vd*, started to develop in ‘Ac-15’ and ‘Picual’ plants, as expected, by 21 and 27 days after pathogen inoculation, respectively (**Supplementary Figure S1**). The mean incubation period was of 21.5±1.7 and 29.5±1.7 days for ‘Ac-15’ and ‘Picual’, respectively. All ‘Picual’ and ‘Ac-15’ plants were dead by 2 months after inoculation. On the contrary, no symptoms developed on nursery and *in vitro*-adapted propagated ‘Ac-18’ plants as expected (Jiménez-Fernández et al., 2016). Surprisingly, plants of ‘Ac-18’ that underwent *in vitro* propagation under aseptic conditions (i.e., *in vitro*-standard) started to develop disease symptoms 29 after inoculation (mean incubation period of 33.8±1.8 days), reaching a disease incidence of 100% and a final disease severity of 1.95±0.6 on a 0-4 rating scale at the end of the experiment (**Figure 1; Supplementary Figure S1**). *Vd* was not re-isolated from any of the stem zones sampled from ‘Ac-18’ nursery and *in vitro*-adapted propagated plants, but was isolated from all ‘Ac-18’ *in vitro*-standard propagated plants, with a mean SCI value of 80.95±6.55%. No symptoms developed on non-inoculated control plants (**Supplementary Figure S1**).

**Figure 1.**
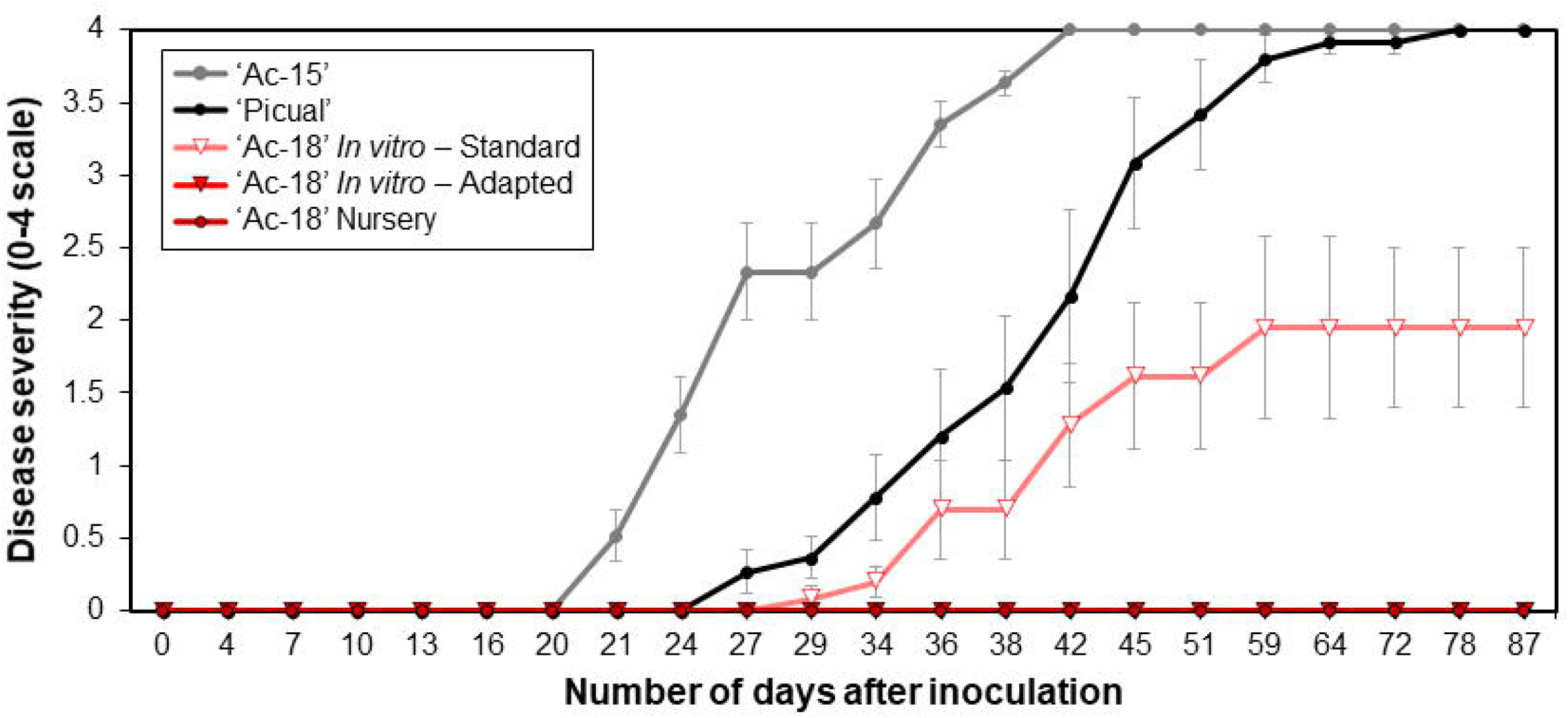
*Verticillium* wilt disease progression in ‘Ac-18’ *in vitro* (standard and adapted) and nursery propagated olive plants infested with the defoliating pathotype of *Verticillium dahliae*. ‘Picual’ and ‘Ac-15’ olive genotypes were used as positive control to determine the inoculation success and development of the disease. Each point represents the mean disease severity (0–4 scale: 0= healthy, 4□=□dead plant) of data and error bars shows the standard error from six plants per treatment.

In parallel with the *Vd* isolations, the pathogen was detected in DNA samples from all *in vitro*-standard propagated plants, with mean Ct values of 28.84±0.18. In contrast, *Vd* was not detected in DNA samples of any of the nursery propagated plants; and only in three samples from the *in vitro*-adapted propagated plants, that showed Ct values slightly above the detection limit (i.e. 37.0±0.30).

### Alpha and beta diversity analysis

Sequencing analysis resulted in a total of 19,936 good quality reads with a mean length of 333 bp after removal of chimeras, unassigned or mitochondrial reads. No chloroplasts reads were detected in our samples. A total of 118 OTUs were identified for all treatments, with 18 OTUs being retained for alpha and beta diversity analysis after rarefying all data to the minimum number of reads obtained and singleton removal. High values of Good’s coverage were obtained for all samples (*data not shown*) indicating enough sequencing depth.

Rarefaction curves of observed OTUs clearly showed a higher number of OTUs on *Vd*-inoculated plants when compared to that on non-inoculated plants, with lower differences among propagation methods within them (**Supplementary Figure S2)**. Similarly, the Scheirer–Ray–Hare test indicated significantly differences (*P* < 0.05) for all alpha-diversity indexes (Richness, Shannon, and Simpson) according to the inoculation treatment (H > 9.94, *P* < 0.002), with no significant differences (*P* ≥ 0.05) among propagation methods (H < 5.14, *P* > 0.076) or its interaction with the inoculation treatment (H < 1.08, *P* > 0.581) (**Figure 2; Supplementary Table S1)**. Interestingly, inoculation with *Vd* significantly increased the number of OTUs identified in all types of propagation (**Figure 1; Supplementary Figure S2**).

**Figure 2.**
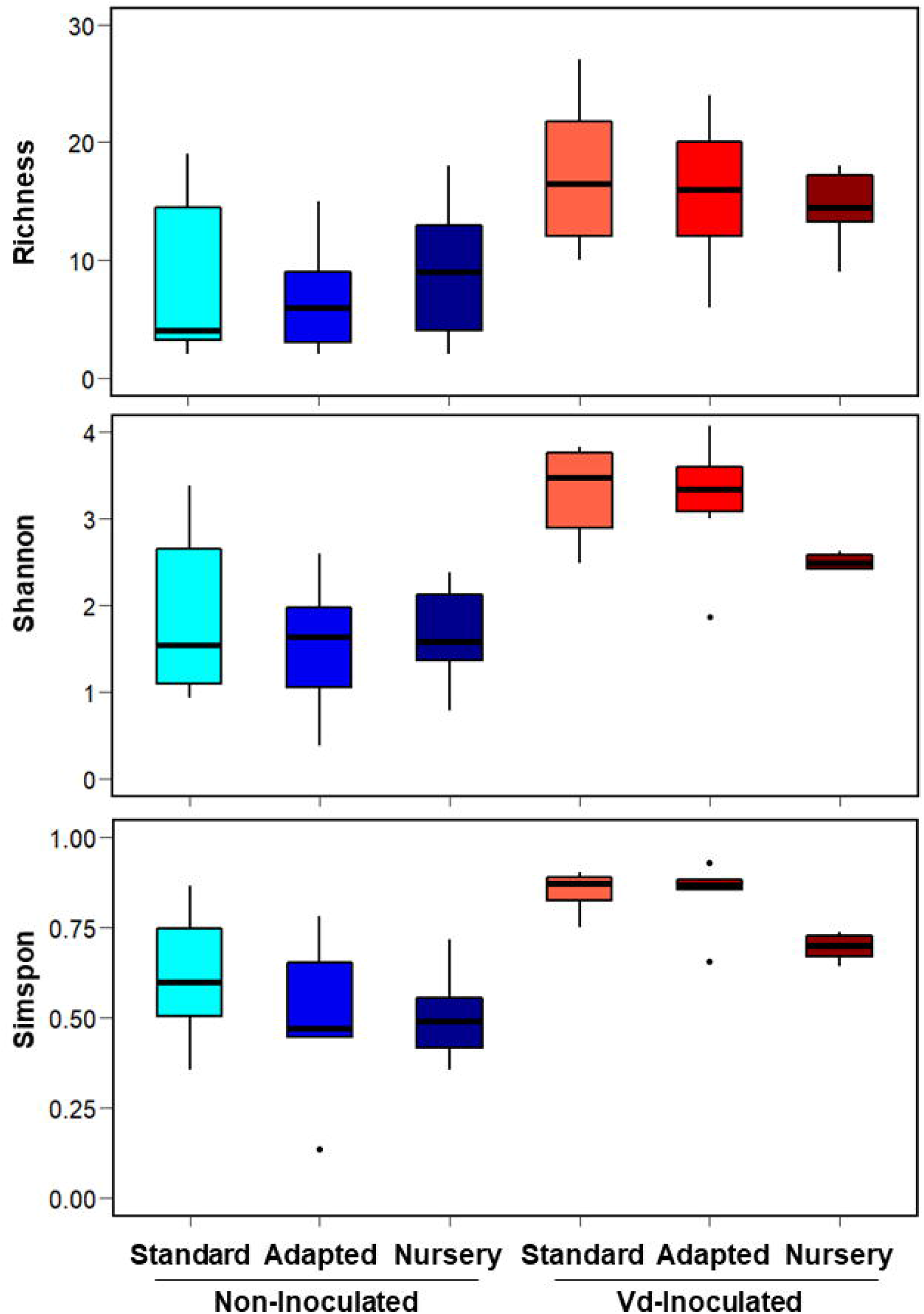
Boxplots of Richness, Simpson and Shannon alpha diversity indices at OTU taxonomic level in olive xylem from *Verticillium dahliae* (*Vd*)-inoculated and non-inoculated ‘Ac-18’ plants following *in vitro* (standard and adapted) and nursery propagation methods. Boxes represent the interquartile range while the horizontal line inside the box defines the median and whiskers represent the lowest and highest values of six values for each treatment combination. For all three indexes and propagation methods, values on *Vd*-inoculated plants were significantly higher compared to that on non-inoculated treatments according to the Scheirer–Ray–Hare test at *P* < 0.05.

PCoA of weighted UniFrac distances differentiated xylem bacterial communities according to the inoculation treatment. Thus, with a few exceptions, there was a clear trend to group the bacterial communities firstly by the presence of *Vd* in the soil mixture, and then according to the propagation method, with *in vitro*-adapted and nursery propagated plants being more similar between them (**Figure 3**). ADONIS analysis indicated a significant effect of the propagation method (*R*^2^ = 0.268, *P* < 0.001) followed by the inoculation treatment (*R*^2^ = 0.112, *P* = 0.004), with no interaction effect (*R*^2^ = 0.062, *P* = 0.175) (**Supplementary Table S2**).

**Figure 3.**
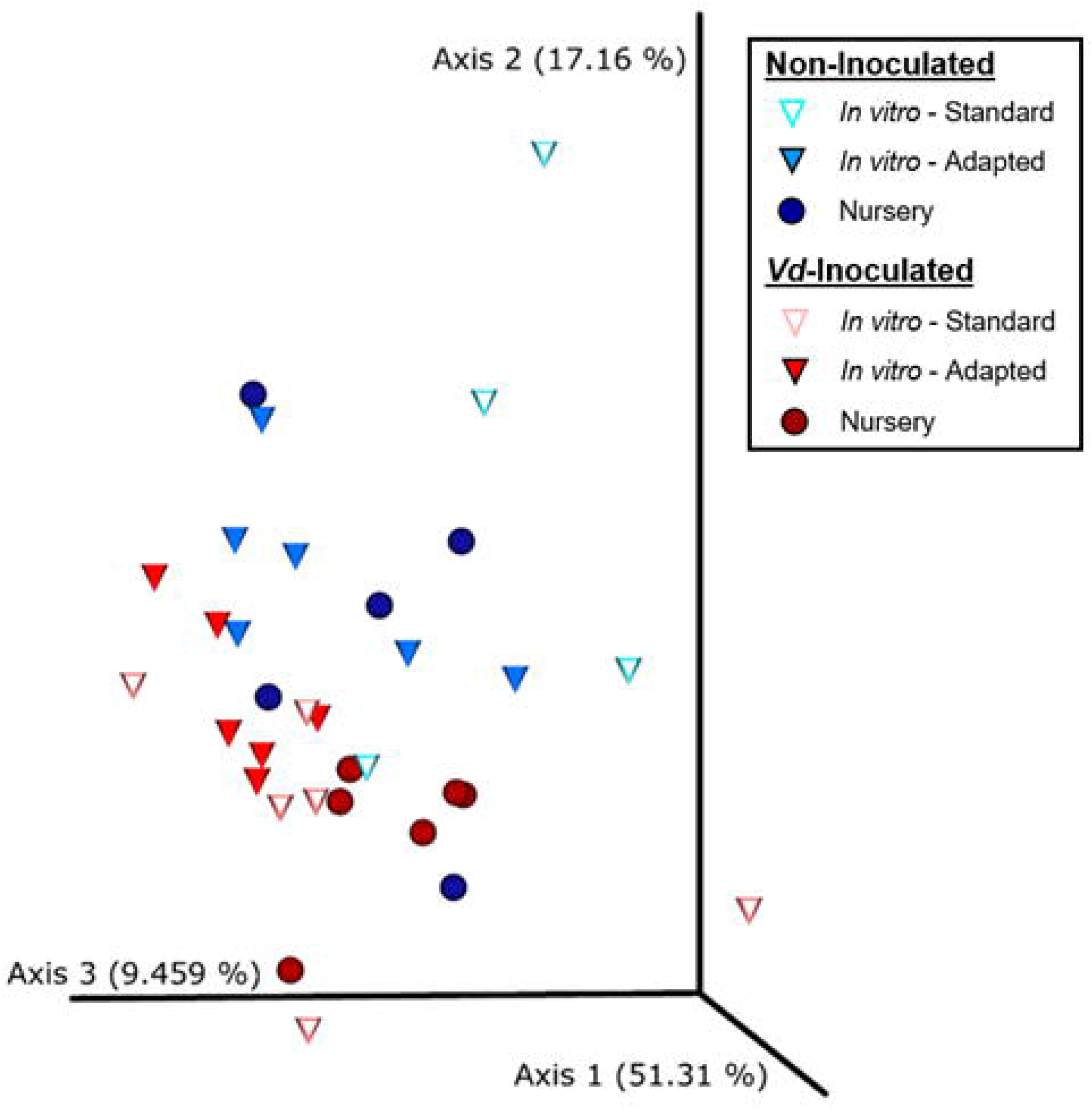
Principal coordinates plot of weighted UniFrac distances of bacterial communities at OTUs taxonomic level in olive xylem from *Verticillium dahliae* (*Vd*)-*inoculated* and non-inoculated ‘Ac-18’ plants following *in vitro* (standard and adapted) and nursery propagation methods. Points are colored by plant inoculation treatment and shaped by propagation methods.

### Composition of olive xylem bacterial communities

A total of 10 phyla, 15 classes, 45 orders, 67 families, 103 genera and 118 OTUs were identified considering all treatments, with four phyla, five classes, 11 orders, 11 families, 10 genera and five OTUs being shared among all of them (**Supplementary Figure S3**). When analyzing the samples according to the inoculation procedure, a higher number of OTUs was found in ‘Ac-18’ plants inoculated with *Vd* (94), while 61 were found on non-inoculated plants. This trend of detecting higher number of taxa on *Vd*-inoculated plants was observed at all the taxonomic levels. Interestingly, the lowest number of OTUs from all taxonomy ranks was found in non-inoculated plants propagated under *in vitro* conditions. Additionally, *in vitro*-adapted plants shared higher number of OTUs with nursery than with *in vitro*-standard propagated plants (**Supplementary Figure S3**).

At genus level, a total of 18 genera formed the core microbiome of *Vd* - inoculated plants when considering all propagation methods jointly, whereas 10 genera were shared within non-inoculated plants. Those same 10 genera were shared between *Vd*-inoculated and non-inoculated treatments (*Acidibacter, Anoxybacillus, Bradyrhizobium, Corynebacterium, Cutibacterium, Methylobacterium-Methylorubrum, Paenibacillus, Pseudomonas, Sphingomonas, Staphylococcus*), while eight genera (*Acinetobacter, Caulobacter, Comamonadaceae, Dermacoccus, Flavisolibacter, Massilia, Paracoccus, Sericytochromatia*) were detected exclusively in *Vd*-inoculated plants. In addition, a higher number of genera was found in *Vd*-inoculated plants (82) compared to those in non-inoculated plants (53) (**Figure 4A**).

**Figure 4.**
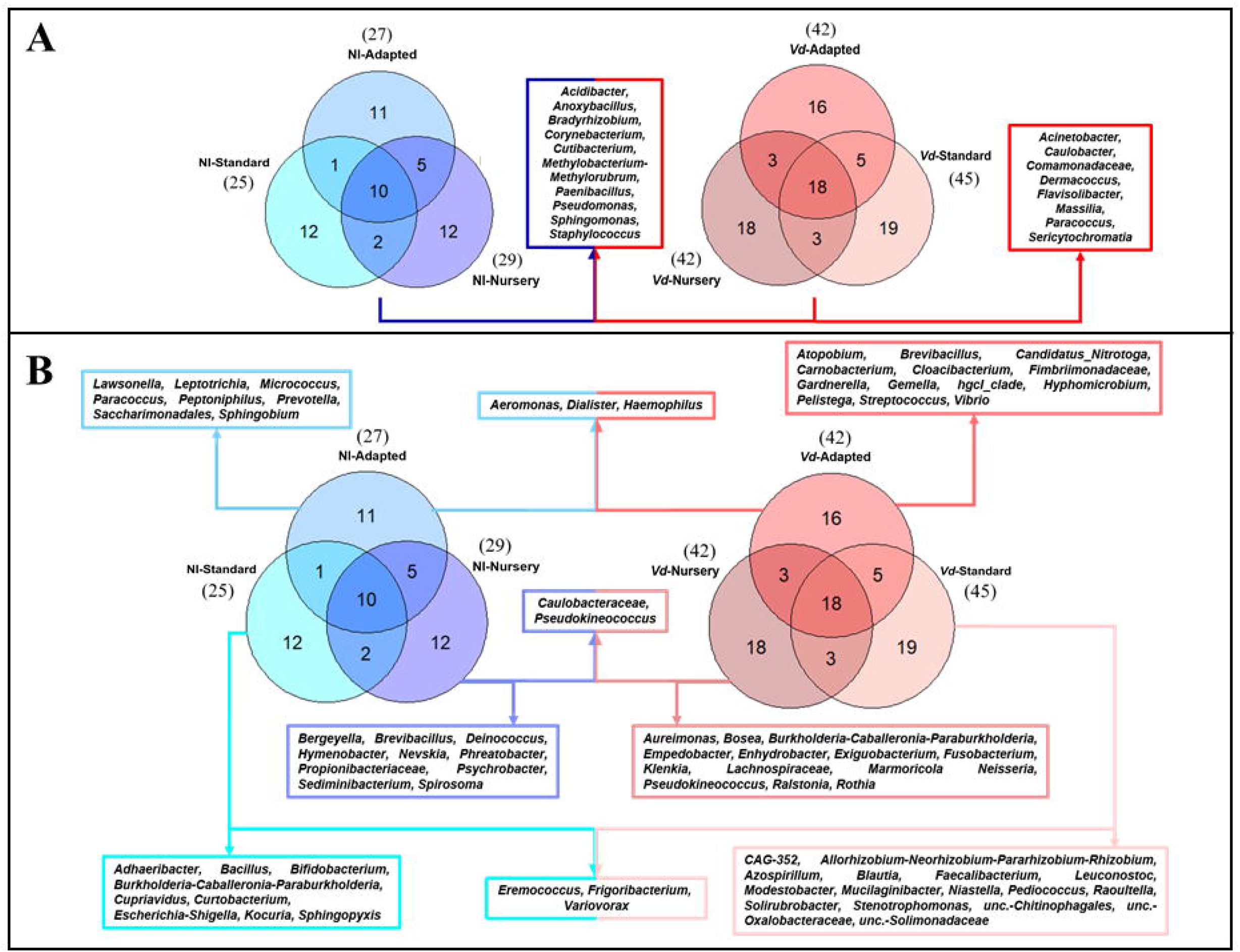
Prevalence Venn diagram showing the unique and shared bacterial genera in olive xylem from the core microbiome (A) or by each propagation approach (B) obtained from *Verticillium dahliae* (*Vd*)-inoculated and non-inoculated ‘Ac-18’ plants following *in vitro* (standard and adapted) and nursery propagation methods.

When analyzing only non-inoculated plants, 25 and 27 genera were identified in *in vitro*-standard and *in vitro*-adapted propagated plants, respectively, while 29 genera were identified in nursery propagated plants. Additionally, a high number of unique genera were found in each propagation method. A total of 12 unique genera were identified for *in vitro*-standard propagated plants (*Adhaeribacter, Bacillus, Bifidobacterium, Burkholderia-Caballeronia-Paraburkholderia, Cupriavidus, Curtobacterium, Eremococcus, Escherichia-Shigella, Frigoribacterium, Kocuria, Sphingopyxis, Variovorax*), 12 for nursery propagated plants (*Bergeyella, Brevibacillus, Caulobacteraceae, Deinococcus, Hymenobacter, Nevskia, Phreatobacter, Propionibacteriaceae, Pseudokineococcus, Psychrobacter, Sediminibacterium, Spirosoma*) and 11 for *in vitro* adapted plants (*Aeromonas, Dialister, Haemophilus, Lawsonella, Leptotrichia, Micrococcus, Paracoccus, Peptoniphilus, Prevotella, Saccharimonadales, Sphingobium*) (**Figure 4B**). On the other hand, when analyzing each propagation methodology for *Vd*-inoculated plants 45, 42 and 42 genera were identified in *in vitro*-standard, *in vitro*-adapted and nursery propagated plants, respectively. Unique genera differed according to the propagation approach. Thus, a total of 19 unique genera were found in *in vitro*-standard propagated plants (*CAG-352, Allorhizobium-Neorhizobium-Pararhizobium-Rhizobium, Azospirillum, Blautia, Eremococcus, Faecalibacterium, Frigoribacterium, Leuconostoc, Modestobacter, Mucilaginibacter, Niastella, Pediococcus, Raoultella, Solirubrobacter, Stenotrophomonas, unc.-Chitinophagales, unc.-Oxalobacteraceae, unc.-Solimonadaceae, Variovorax*), 16 genera were exclusive from *in vitro*-adapted propagated plants (*Aeromonas, Atopobium, Brevibacillus, Candidatus_Nitrotoga, Carnobacterium, Cloacibacterium, Dialister, Fimbriimonadaceae, Gardnerella, Gemella, Haemophilus, hgcI_clade, Hyphomicrobium, Pelistega, Streptococcus, Vibrio*) and 18 from nursery propagated plants (*Actinomyces, Aeromicrobium, Agromyces, Aureimonas, Bosea, Burkholderia-Caballeronia-Paraburkholderia, Caulobacteraceae, Empedobacter, Enhydrobacter, Exiguobacterium, Fusobacterium, Klenkia, Lachnospiraceae, Marmoricola Neisseria, Pseudokineococcus, Ralstonia, Rothia*) (**Figure 4B**).

### Bacterial abundance

At phylum level, *Actinobacteriota* presented the highest relative abundance considering all experimental treatments together (43.62%), followed by *Proteobacteria* (38.72%), *Firmicutes* (15.24%) and *Bacteroidota* (1.63%). However, these relative abundances varied within each treatment tested. *Actinobacteria* was more abundant in non-inoculated plants, with a proportion of 46.41%, decreasing to 40.82% for *Vd*-inoculated plants. For non-inoculated plants, this Phylum was the most abundant for *in vitro*-adapted propagated plants (70.98%), followed by nursery (52.21%) and *in vitro*-standard (16.04%) propagation methods; showing the same trend in *Vd*-inoculated plants, in which reached 58.56, 39.16 and 24.73%, for these same propagation methods, respectively. *Proteobacteria* were present at similar percentage in non-inoculated (40.77%) and *Vd*-inoculated (36.66%) plants, but varied within propagation methods. Thus, it was the most abundant phylum for the *in vitro*-standard plants both for non-inoculated (75.86%) and *Vd*-inoculated plants (49.94%), being lowest for *in vitro*-adapted propagated plants with 18.79 and 22.12%, for non- and *Vd*-inoculated plants, respectively. Finally, *Firmicutes*, showed a different response to both, inoculation treatment and propagation methods compared to the two previous Phyla. Firstly, it showed the highest global abundance in *Vd*-inoculated plants with 18.92%, that decreased to 11.56% in non-inoculated plants. Secondly, while in non-inoculated plants, the highest abundance was estimated in nursery propagated plants (17.53%), it represented ca. 8.57% in both *in vitro* propagation methods; but similar abundance values were reached in *Vd*-inoculated plants (17.61 to 20.34%), irrespective of the plant propagation method (**Figure 5A**).

**Figure 5.**
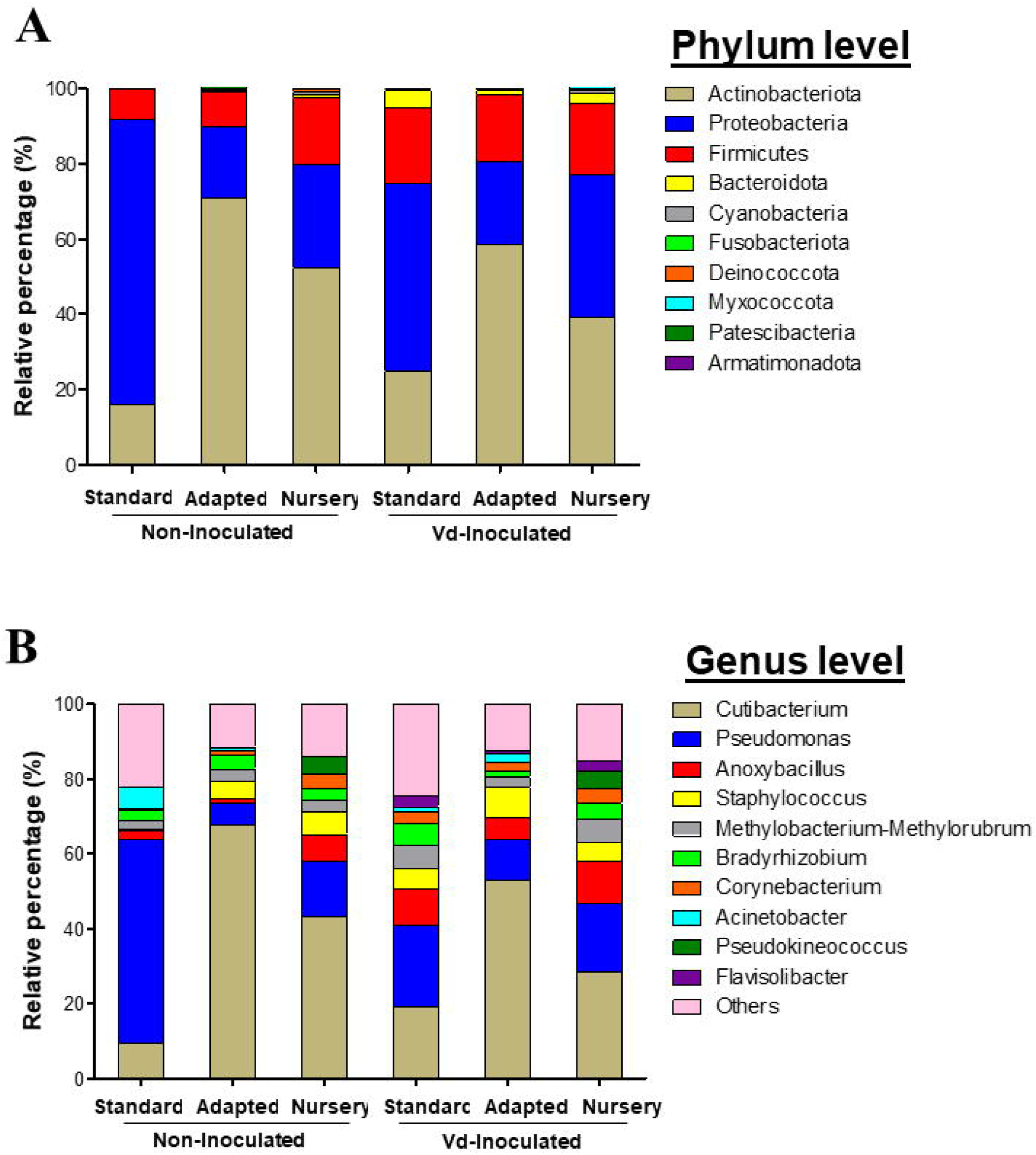
Barplots showing the relative bacterial abundance taxa at phylum (A) and genera (B) level present in olive xylem from *Verticillium dahliae* (*Vd*)-inoculated and non-inoculated ‘Ac-18’ plants following *in vitro* (standard and adapted) and nursery propagation methods.

In line with these results, Least discriminant analysis effect size (LEfSe) was used to identify the key phylotypes that could be differentially associated to the different experimental treatments. When comparing the three propagation methods within each inoculation treatment, Proteobacteria-Gammaproteobacteria was the most significant Phylum-Class for non-inoculated *in vitro*-standard propagated plants. For plants growing in *Vd*-infested soils, more diversity was observed among propagation methods, with Proteobacteria-Gammaproteobacteria and Bacteroidota-Bacteroidia, and Actinobacteriota-Actinobacteria being the most prevalent Phylum-Class in *in vitro*-standard and in *in vitro*-adapted propagated plants, respectively, while the Class-Family Proteobacteria-Alphaproteobacteria was the most prominent in nursery propagated plants. On the other hand, when comparing the effect of the inoculation with the pathogen within each propagation method, Firmicutes-Bacilli was a significant Phylum-Class for *Vd*-inoculated plants, in both, *in vitro*-standard and *in vitro*-adapted propagated plants, whereas Proteobacteria-Gammaproteobacteria and Bacteroidota-Bacteroidia were also the prevalent Phyla-Class for *in vitro*-standard and nursery propagated plants, respectively. For non-inoculated plants only a Phylum-Class (Actinobacteriota-Actinobacteria) appeared as the most prominent and only for *in vitro*-adapted plants (**Figure 6**).

**Figure 6.**
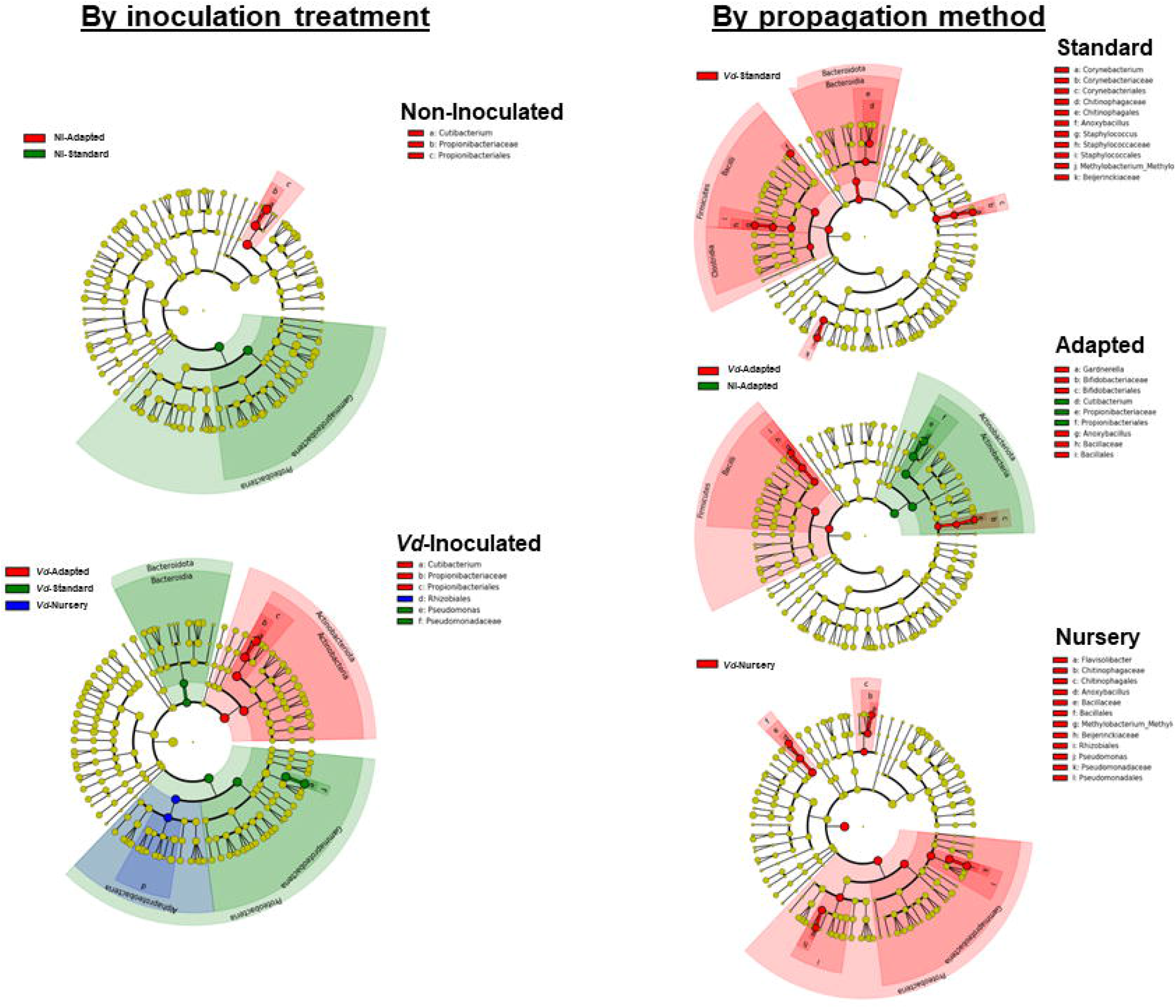
Cladogram representation from LEfSe analysis showing the taxonomic ranks from innermost phylum ring to outermost genera ring. Each point is a member within each taxonomic rank. Significant taxa (*P* < 0.05) by inoculation treatment (Vd: *Verticillium dahliae* (*Vd*)-inoculated; NI: non-inoculated plants) or by propagation method (*in vitro* standard and adapted, and nursery propagated plants) are shown in different colors.

The most abundant genera identified among all treatments were *Cutibacterium* (36.85%), *Pseudomonas* (20.93%), *Anoxybacillus* (6.28%), *Staphylococcus* (4.95%), *Methylobacterium-Methylorubrum* (3.91%), *Bradyrhizobium* (3.54%), *Corynebacterium* (2.53%), *Acinetobacter* (1.77%), *Pseudokineococcus* (1.59%) and *Flavisolibacter* (1.07%) (**Figure 5B**). *Cutibacterium* was the genus with the highest relative abundance, reaching maximum frequencies in *in vitro*-adapted plants [both in non-inoculated (67.81%), and *Vd*-inoculated plants (53.04%)]. Other predominant genera varied in proportion depending upon the treatment combination. *Pseudomonas* (54.25%) was the most representative genera in non-inoculated plants, propagated by the *in vitro*-standard method, whereas similar abundances were found for the remaining treatments (14.66%), with the exception of *in vitro*-adapted non-inoculated plants (5.77%). A noticeable lower abundance of *Anoxybacillus* was found in non-inoculated plants from both *in vitro* propagation methods (standard and adapted with 2.42% and 1.06%, respectively) compared with the nursery propagation (7.32%). Also, it was remarkable the small proportion of *Staphylococcus* found in *in vitro*-standard propagated and non-inoculated plants (0.52%) compared with the rest of the treatments (5.84%). Also, *Acinetobacter* showed a high proportion (3.34%) in *in vitro* propagated plants, but were much less abundant in nursery propagated plants (0.04%) (**Figure 5B**).

In line with these results, when comparing the propagation methods within each inoculation treatment, LEfSe displayed *Cutibacterium* as the only significant genus when plants were not inoculated, but only for *in vitro*-adapted propagated plants; whereas for *Vd*-inoculated plants *Anoxybacillus* appeared as a significant genus for all propagation methods (**Figure 6**). Furthermore, other distinct genera appeared as the most dominant within each propagation method when *Vd* was infesting the soil including *Flavisolibacter, Methylobacterium-Methylorubrum*, and *Pseudomonas* in nursery propagated plants; *Corynebacterium, Staphylococcus* and *Methylobacterium-Methylorubrum* in *in vitro*-standard propagated plants and *Gardnerella* in *in vitro*-adapted plants. Finally, when comparing the three propagation methods *Pseudomonas* appeared as the most dominant genera for *in vitro*-standard propagated plants infected by *Vd* (**Figure 6**).

## Discussion

Olive is a crop of utmost ecological, economic and cultural significance which must be preserved for next generations (Besnard et al., 2013). However, old and emerging xylem-inhabiting pathogens, such as *Vd* or *Xylella fastidiosa* endanger its health status nowadays (Jiménez-Díaz et al., 2011; Schneider et al., 2020). The use of host resistance is the most practical, long-term and economically efficient disease control measure for Verticillium wilt in olive, and it is at the core of integrated disease strategies that must be practiced for the efficient management of the disease. Use of wild olive rootstocks highly resistant to D *Vd* can provide an improved means for the management of Verticillium wilt, especially for grafting susceptible olive cultivars that are agronomically adapted, commercially desirable or used in protected designation of origin extra virgin olive oils (Jiménez-Díaz et al., 2011; Trapero et al., 2012, 2013; Jiménez-Fernández et al., 2016; Ostos et al., 2020). In addition to that, use of endophytic plant-associated microorganisms with a specific beneficial interaction with the host plant could help to improve olive health and productivity providing a potential perspective for sustainable plant protection (Ryan et al., 2008; Berg, 2009; Berg et al., 2014; Müller et al., 2015).

In olive, several defense mechanisms, including both biochemical responses and plant structural characteristics, have been proposed as factors contributing to the resistance shown by different genotypes against its vascular pathogens *Vd* and/or *X. fastidiosa*. Those mechanisms should operate within the xylem tissues contributing to reduce systemic colonization by the pathogen, and may include build-up of vessel occlusions by gums, gels or tyloses, phenolics and lignin content or their accumulation, induction of pathogenesis-related proteins, antioxidant-related enzymes, and ionome content (Báidez et al., 2007; Markakis et al., 2010; Jiménez-Fernández et al., 2016; Gharbi et al., 2017a, 2017b; Leyva-Pérez et al., 2017; Luvisi et al., 2017; Sabella et al., 2017, 2019; D’Attoma et al., 2019).

Although some cultivated and wild olive clones, including the ‘Ac-18’ used in this study, have been described as resistant to D *Vd* based on symptomless resistance; for most of them the pathogen could be detected by molecular methods or re-isolated from stem tissues (Bubici and Cirulli, 2012; Colella et al., 2008; Gramaje et al., 2013; Jiménez-Fernández et al., 2016) indicating the plant’s ability to reduce the extent of stem colonization or other pathogenesis mechanisms that result in the absence of visible disease symptoms. However, the role that xylem microbial communities may play in that resistant response to *Vd* has been overlooked and remains unexplored to date. In this study, we tested the hypothesis that xylem microbiome may have a functional role on plant resistance. With that purpose, we explored whether or not *in vitro* propagation of ‘Ac-18’ plants can alter the diversity and composition of xylem-inhabiting bacteria, and to which extent this could result in a modification of the high resistance response of that wild olive clone to the highly virulent D pathotype of *Vd*. Surprisingly, plants that underwent *in vitro* propagation under aseptic conditions lost the high resistance phenotype characteristic of the ‘Ac-18’ clone. Actually, those plants developed wilting symptoms similar to those reported for other olive cultivars with a moderate-susceptible reaction to D *Vd* such as ‘Frantoio’ ‘Oblonga’ ‘Koroneiki’ ‘Empeltre’ or ‘Leccino’ in similar inoculation experiments using olive plants of age similar to that in our study (López-Escudero et al., 2004; Martos-Moreno et al., 2006; Trapero et al., 2013).

Resistance to fungal vascular wilts may change during plant growth and development. Some authors found that disease severity in olive cultivars susceptible to *Vd* decreases with host age (López-Escudero et al., 2010; Trapero et al., 2013). In this present study, the loss of resistance shown by ‘Ac-18’ *in vitro*-standard propagated plants cannot be associated to a more juvenile stage since those plants were of the same age than that of nursery-propagated plants, and showed a similar growth (i.e., similar bark lignification and root development). However, ‘Ac-18’ plants showed distinct xylem microbiome profiles according to the propagation procedure. The most significant change associated to *in vitro*-standard propagation was a decrease in the total number of OTUs detected, and a significantly higher number of Gammaproteobacteria (mainly *Pseudomonas*) and lower number of Actinobacteria (mainly *Cutibacterium*). In parallel, alpha- and beta-diversity indexes of xylem microbiome differed among propagation procedures, with plants that were initially propagated under *in vitro*-standard conditions and then challenged to a less restricted aseptic environmental conditions (i.e., *in vitro*-adapted plants) showing a xylem microbiome more similar with the commercial nursery propagated plants. The olive explants from tissue culture may contain a genotype-specific core xylem microbiome that is transmitted from shoot tips of last generation (Liu et al., 2019). In our study, the explants grew under aseptic conditions and roots that differentiated from them did not contact with outside microbes, at least until the challenge with the pathogen. Thus, most of the differences found in *in vitro*-adapted or nursery propagated plants may be attributed to bacteria that were present at very low level, bellow the detection limit, at the beginning of micropropagation procedure and could not be detected by NGS. Alternatively, those bacteria might have been acquired by roots after recruitment when plants grew on a non-sterile substrate under less aseptic environmental conditions as proposed for other woody crops, including olive (Antoniou et al., 2017; Fausto et al., 2018; Deyett and Rolshausen, 2019).

Plant core microorganism are considered to be consistently established in plants not being influenced by differences in space, time or plant organs (Vorholt et al., 2017). In our study, 10 keystone bacterial genera could be considered the core microbiome being transmitted from generation to generation in olive, since they were detected in all samples regardless plant propagation procedure (*in vitro* vs nursery) or inoculation with the pathogen; with *Cutibacterium, Pseudomonas, Anoxybacillus, Staphylococcus, Methylobacterium-Methylorubrum*, and *Bradyrhizobium* being the most abundant, in that order. These bacterial taxa have also been identified in olive xylem in other works in which olive trees of different ages, belonging to different cultivars or growing under different environments were evaluated (Müller et al. 2015; Fausto et al., 2018; Sofo et al., 2019; Anguita-Maeso et al., 2020; Giampetruzzi et al., 2020) which strength the hypothesis that those genera may represent keystone olive xylem bacteria. More interestingly, some of these genera have been already reported or used as plant growth promoting bacteria (Otieno et al., 2015; Subramanian et al., 2015) or as biological control agents against *V. dahliae* (Berg et al., 2002, 2006). Interestingly, the ratio *Cutibacterium/Pseudomonas* seemed to be an important factor associated to the plant propagation procedure. Thus, *Pseudomonas* spp. and *Cutibacterium* were present at high and low abundance, respectively, for *in vitro*-standard propagated plants, that lost resistance to D *Vd*, whereas for *in vitro*-adapted and nursery propagated plants the opposite trend occurred. However, little is known about the role of the genus *Cutibacterium* as a component of plant microbiome, whereas the beneficial functions of *Pseudomonas* spp. in plants have been widely reported for several crops, including olive (Mercado-Blanco et al., 2004; Weller, 2007; Loper et al., 2012).

The role of microorganisms in the biocontrol of Verticillium wilt diseases has been reported mostly on non-woody plant species such as cotton, potato, strawberry or tomato (Azad et al., 1985; Nallanchakravarthula et al., 2014; Cao et al., 2016; Wei et al., 2019; Snelders et al., 2020) with few studies focused on woody hosts including olive (Mercado-Blanco et al., 2004; Aranda et al., 2011; Gómez-Lama Cabanás et al., 2018). However, the characterization of microbial communities inhabiting xylem vessels colonized by *Vd* has not been studied to date, despite some work done on other tree species or other vascular pathogens (Martín et al., 2015; Pérez-Martínez et al., 2018; Giampetruzzi et al., 2020; Vergine et al., 2020). To the best of our knowledge, this present study is the first to address this knowledge gap, by determining changes in xylem bacterial communities of a resistant olive clone after challenge inoculation with D *Vd*. Our results indicated that a significantly higher diversity and number of OTUs occurs in *Vd*-inoculated plants regardless of the plant propagation method and success of stem vascular infection by the pathogen. Additionally, several genera appeared as unique in *Vd*-inoculated plants including *Acitenobacter, Caulobacter*, an uncultured Comamonadaceaeae, *Dermacoccus, Flavisobacter, Massilia, Paracoccus* and *Sericytochromatia*. Also, *Anoxybacillus;* represented a keystone bacterial taxa that significantly increased its frequency after challenge inoculation with the pathogen in all treatments.

The significantly higher xylem microbial diversity in *Vd*-inoculated plants is in line with results from other studies involving vascular pathogens on woody crops. For instance, Deyett and Rolshausen (2019) found higher diversity in *X. fastidiosa-infected* vines as compared to healthier ones. Several of the unique or keystone bacterial genera (e.g., *Acinetobacter, Comamonas, Caulobacter, Massilia, Methylobacterium*) detected in our study in the xylem of *Vd* inoculated plants were also found in the xylem of other plant species such as banana, citrus, grapevine and olive, so that those bacteria may be biomarkers of plant infection by vascular-plant pathogens (Araújo et al., 2002; Deyett and Rolshausen, 2019; Liu et al., 2019). The role of these bacterial genera in *Vd*-infected olive plants remains unknown, although several studies suggested that the antifungal activity of *Acinetobacter* or *Dermacoccus* might be involved in pathogen suppression (Liu et al., 2007; AlMatar et al., 2017). *Methylobacterium* isolates have been shown to induce resistance against attack by diverse plant-pathogenic fungi (Rajendran et al., 2009; Ardanov et al., 2012) and modify the response of citrus to the xylem-inhabiting bacterial pathogen *X. fastidiosa*, resulting in a reduction of Citrus Variegated Chlorosis symptoms (Azevedo et al., 2016). *Anoxybacillus*, a keystone bacteria that increased its abundance in pathogen inoculated-plants, can produce several cellulose-, hemicellulose-, lignin- and starch-degrading enzymes, whose activity might have been induced following infection by *Vd* in olive vascular tissue, thus facilitating its population increase (Goh et al., 2013).

The general increase in alpha diversity and abundance of specific bacterial taxa observed in our study after challenging with D *Vd* may be explained by several hypotheses, including: i) a passive entry or direct recruitment of new bacterial species from the plant rhizosphere or soil, taking advantage of injuries caused during root infection and colonization by the pathogen (Ardanov et al., 2012; Liu et al., 2019); ii) the secretion of specific molecules by the pathogen (such as effector proteins) with antimicrobial activity that modify host microbiome to facilitate host colonization (Snelders et al., 2020); and/or iii) the pathogen provokes a series of host physiological responses that trigger multiplication of a specific microbiome to cope with the pathogen infection in order to mitigate its effect (Ardanov et al., 2012; Hassani et al., 2018; Carrión et al., 2019). These hypotheses emphasize the need for better understanding of the changes occurring in xylem microbial communities in response to vascular infection by pathogens, in order to determine specifically activated disease-suppressive and/or plant-protecting microbiome-mediated activities in olive.

Our study provides new insights for the characterization of changes occurring in the xylem microbial communities of a wild olive genotype following inoculation with the vascular plant pathogen *Vd*. Also, it provides a quantitative and qualitative assessment of the effect of specific propagation methods where the attenuation and reduction of olive xylem microbiome is a unique approach described to date. We are aware of limitations in our study, in part because only some propagation methods and a single host genotype were evaluated. However, this present research is relevant for future studies on the olive xylem microbiome that may lead to identification of xylem-inhabiting bacteria potentially involved in host resistance and plant defense by acting as biocontrol agents against xylem-inhabiting pathogens. Deciphering the core olive xylem microbiome and their correlation with the host plant and its pathogens is a first critical step for exploiting the microbiome in order to enhance olive growth and health.

## Author Contributions

MA-M and BBL: conceived research, performed statistical and bioinformatics analyses, interpret results and wrote the manuscript; MA-M, JLT, CO-G, DR-R and EP-R: prepare materials and equipment and performed the experiments; JAN-C and RMJ-D: contributed to reviewing the manuscript and interpret results. All authors viewed the draft of manuscript.

## Acknowledgments

We acknowledge support of the publication fee by the CSIC Open Access Publication Support Initiative through its Unit of Information Resources for Research (URICI).

## Data Availability Statement

The raw sequence data have been deposited in the Sequence Read Archive (SRA) database at the NCBI under BioProject accession number PRJNA679263.

## Supplementary Material

**Figure S1.** Effect of inoculation with the defoliating pathotype of *Verticillium dahliae* in ‘Picual’, ‘Ac-15’ and ‘Ac-18’, plants obtained using *in vitro*-standard, *in vitro*-adapted and nursery propagation methods. ‘Picual’ and ‘Ac-15’ were used as positive control to determine the inoculation success and the development of the disease. Note the defoliation of green leaves observed for *in vitro*-standard propagated ‘Ac-18’ plants.

**Figure S2.** Richness rarefaction curves at OTU taxonomic level in olive xylem from *Verticillium dahliae* (*Vd*)-inoculated and non-inoculated (NI) ‘Ac-18’ plants following *in vitro* (standard and adapted) and nursery propagation methods. Error bars represent standard error of six values.

**Figure S3**. General prevalence Venn diagram showing the unique and shared bacteria at different taxonomy ranks in olive xylem from *Verticillium dahliae* (*Vd*)-inoculated and non-inoculated (NI) ‘Ac-18’ plants following *in vitro* (standard and adapted) and nursery propagation methods.

