## Supplementary figures and images for "*Verticillium dahliae* inoculation and *in vitro* propagation modify the xylem microbiome and disease reaction to Verticillium wilt in a wild olive genotype"

### Suppl Figure S1

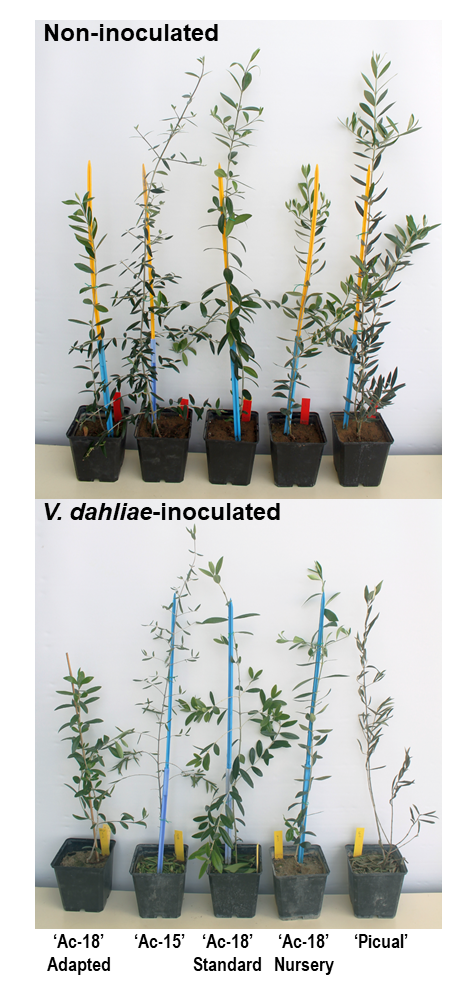

### Suppl Figure S2

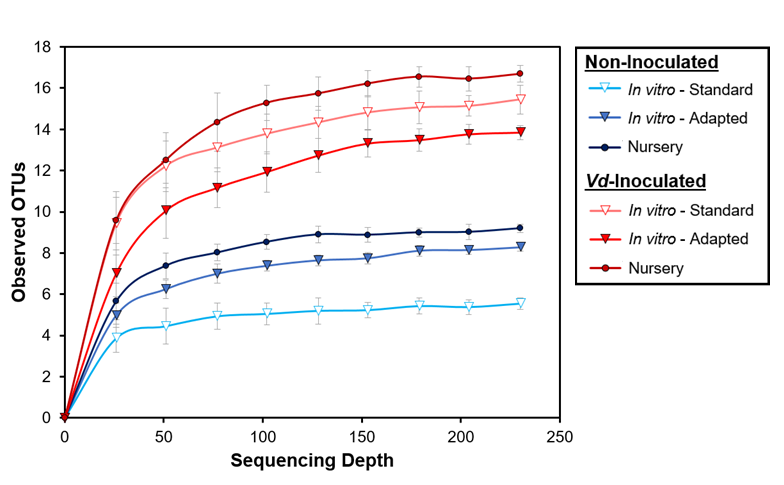

### Suppl Figure S3

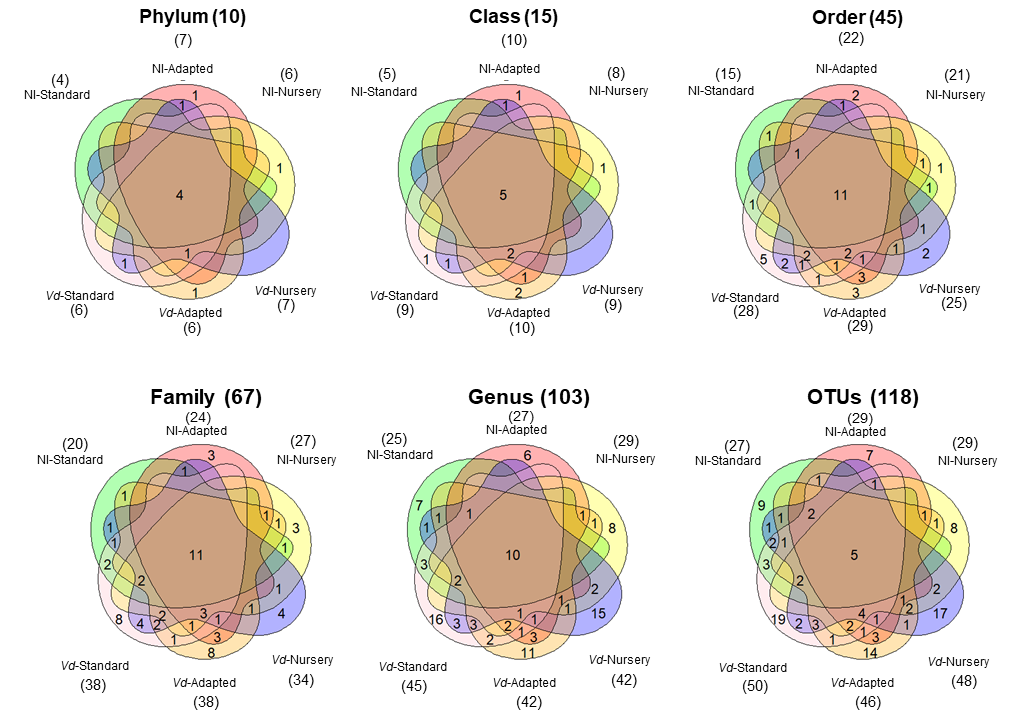
