## Supplementary material for "*Verticillium dahliae* inoculation and *in vitro* propagation modify the xylem microbiome and disease reaction to Verticillium wilt in a wild olive genotype": Suppl Tables

**Table S1.** The Scheirer–Ray–Hare test showing H and *P*-value on Richness, Shannon, and Simpson alpha diversity indices to assess the effects of the inoculation treatment, the plant propagation method (*in vitro-*standard, *in vitro-*adapted and nursery) and their interaction.

| **Diversity index** | **Treatment** | **df** | **Sum of squares** | **H** | ***P*-value** |
| --- | --- | --- | --- | --- | --- |
| Richness | Inoculation treatment | 1 | 1032.27 | 9.903 | **0.0017** |
|  | Propagation method | 2 | 95.82 | 0.919 | 0.6315 |
|  | Inoculation x propagation | 2 | 57.54 | 0.552 | 0.7588 |
|  | Residuals | 29 | 2358.37 |  |  |
| Shannon | Inoculation treatment | 1 | 1674.62 | 15.949 | **< 0.0001** |
|  | Propagation method | 2 | 322.04 | 3.067 | 0.2158 |
|  | Inoculation x propagation | 2 | 90.87 | 0.865 | 0.6488 |
|  | Residuals | 29 | 1482.47 |  |  |
| Simpson | Inoculation treatment (A) | 1 | 1512.66 | 14.406 | **0.0002** |
|  | Propagation method (B) | 2 | 540.20 | 5.145 | 0.0764 |
|  | A x B | 2 | 113.77 | 1.084 | 0.5817 |
|  | Residuals | 29 | 1403.37 |  |  |

**Table S2.** ADONIS test showing *R*^2^ and *P*-values to assess the significance of the inoculation treatment and the plant propagation method and their interaction

| **Treatment** | **df** | **Sequential Sums**  **of Squares** | **Mean Squares** | ***F*** | **partial *R*^2^** | ***P*-value** |
| --- | --- | --- | --- | --- | --- | --- |
| Inoculation treatment | 1 | 0.0514 | 0.0514 | 5.3911 | 0.1115 | **0.004** |
| Propagation method | 2 | 0.1235 | 0.0617 | 6.4780 | 0.2680 | **0.001** |
| Inoculation x Propagation | 2 | 0.0286 | 0.0143 | 1.5008 | 0.0621 | 0.175 |
| Residuals | 27 | 0.2573 | 0.0095 |  | 0.5584 |  |
| Total | 32 | 0.4610 |  |  | 1 |  |
